# Local adaptation and sexual selection drive intra-island diversification in a songbird lineage: differential divergence in autosomes and sex chromosomes

**DOI:** 10.1101/353771

**Authors:** Yann XC Bourgeois, Joris AM Bertrand, Boris Delahaie, Hélène Holota, Christophe Thébaud, Borja Milá

## Abstract

Recently diverged taxa showing marked phenotypic and ecological diversity are optimal systems to test the relative importance of two major evolutionary mechanisms, adaptation to local ecological conditions by natural selection, or mechanisms of reproductive isolation such as assortative mating mediated by sexually selected mating signals or post-zygotic incompatibilities. Whereas local adaptation is expected to affect many loci throughout the genome, traits acting as mating signals are expected to be located on sex chromosomes and have a simple genetic basis. We used genome-wide markers to test these predictions in Reunion Island’s gray-white eye (*Zosterops borbonicus*), which has recently diversified into five distinct plumage forms. Two of them correspond to a polymorphic highland population that is separated by a steep ecological gradient from three distinct lowland forms that show narrow contact zones in plumage color traits, yet no association with environmental variables. An analysis of population structure using genome-wide SNP loci revealed two major clades corresponding to highland and lowland forms, respectively, with the latter separated further into three independent lineages corresponding to plumage forms. Coalescent tests of alternative demographic scenarios provided support for divergence of highland and lowland lineages with an intensification of gene flow in the last 60,000 years. Landscapes of genomic variation revealed that signatures of selection associated with elevation are found at multiple regions across the genome, whereas most loci associated with the lowland forms are located on the Z sex chromosome. A gene ontology analysis identified *TYRP1*, a Z-linked color gene, as a likely candidate locus underlying color variation among lowland forms. Our results are consistent with the role of natural selection in driving the divergence of locally adapted highland populations, and the role of sexual selection in differentiating lowland forms through reproductive isolation mechanisms, showing that both modes of lineage divergence can take place at very small geographic scales in birds.

## Introduction

As populations and lineages diverge from each other, a progressive loss of shared polymorphisms and accumulation of fixed alleles is expected. This is impacted by neutral processes (e.g. genetic drift), but also natural and sexual selection. The interaction between these processes may vary between different parts of the genome, creating a mosaic pattern of regions displaying different rates of divergence(Wu 2001; Nosil et al. 2009). Consequently, regions directly involved in local adaptation and reproductive isolation will experience reduced effective gene flow compared to genomic background (Ravinet et al. 2017). Establishing how different processes such as drift, selection and gene flow, shape the rates of divergence at the genomic scale is critical to understand the speciation process.

Reproductive isolation between nascent species can arise as incompatibilities between interacting loci accumulate in the genome, as described by the Bateson-Dobzhansky-Muller model (Dobzhansky 1936; Muller 1940, 1942; Orr 1996). An excess of highly differentiated regions on sex chromosomes at the early stages of speciation suggests a role for intrinsic barriers to gene flow maintaining divergence (Backström et al. 2010). Sex chromosomes are particularly susceptible to the gathering of incompatibilities, since the genes they harbour are not transmitted by the same rules between males and females (Seehausen et al. 2014) and are therefore prone to genetic conflicts. Indeed, sex chromosomes often harbor many genes causing disruption of fertility or lower viability in hybrids (Carling and Brumfield 2008; Macholán et al. 2011; Ellegren et al. 2012), and are associated with faster emergence of postzygotic isolation (Lima 2014). This large effect of sexual chromosomes has been a common explanation of Haldane’s rule (Haldane 1922; Orr 1997; Coyne and Orr 2004), which states that in hybrids, the heterogametic sex often displays a stronger reduction in fitness.

In addition, premating and prezygotic isolation often affect sexually dimorphic traits that are under the control of sex-linked genes (Pryke 2010). While this type of isolation would lead to divergence along the whole genome due to the isolation of gene pools, divergence is expected to be accelerated at loci controlling traits under sexual selection, especially if hybrids and backcrosses have lower mating success (Svedin et al. 2008). At last, effective recombination rates are lower in sex chromosomes since they recombine in only one sex (males in birds), which facilitates linkage of genes involved in pre- and post-zygotic barriers. This linkage would ultimately promote reinforcement of isolation between species or populations (Pryke 2010). Because of these processes, gene flow at isolating sex-linked loci is expected to be impeded in diverging populations, leaving a stronger signature of differentiation than in the rest of the genome.

On the other hand, adaptation to local environmental conditions or natural counterselection of hybrids is more likely to be driven by genes scattered along the genome (Qvarnström and Bailey 2009; Seehausen et al. 2014). On later stages of the speciation process, coupling between incompatibilities and loci linked to local adaptation can facilitate isolation, and promote reinforcement (Maan and Seehausen 2011; Seehausen et al. 2014; Wolf and Ellegren 2016), even in an initial context of relatively high, continuous gene flow (Kulmuni and Westram 2017). Identifying the main drivers of genome-wide differentiation (i.e. isolation by environment v. reproductive isolation) remains however a complex question (Wolf and Ellegren 2016; Ravinet et al. 2017). Current studies in the field have displayed varied results, and those focusing on the early stages of speciation have often put the emphasis on ecological divergence instead of sexual selection and intrinsic incompatibilities (Bierne et al. 2011; Seehausen et al. 2014). Studies of closely related taxa or populations that show phenotypic and ecological diversity and are at different stages of divergence hold promise to help clarify the chronology and relative importance of these underlying evolutionary mechanisms (Pryke 2010; Sætre and Sæether 2010; Seehausen et al. 2014).

We used the Réunion Grey White-Eye (*Zosterops borbonicus*) to test the relative impacts of ecology and reproductive isolation on genome-wide patterns of divergence on a set of closely related but phenotypically distinct parapatric colour forms. This small passerine endemic to the island of Réunion displays a striking pattern of conspicuous phenotypic variation at a small geographical scale (Figure 1). Three parapatric colour forms are found at low altitude, with a completely brown form (LBHB), a grey-headed brown form (GHB) and a grey headed brown form with a brown nape (BNB). A fourth, polymorphic form is found at high elevation (up to 2,500m), where two colour morphs (brown and grey, HBHB and GRY) coexist without any evidence of genetic structure apart for a single locus on chromosome 1 (Bourgeois et al. 2017). In addition, patterns of colouration among forms and morphs are stable over time, with no sex effect (see Gill 1973, Mila et al. 2010).

**Figure 1.**
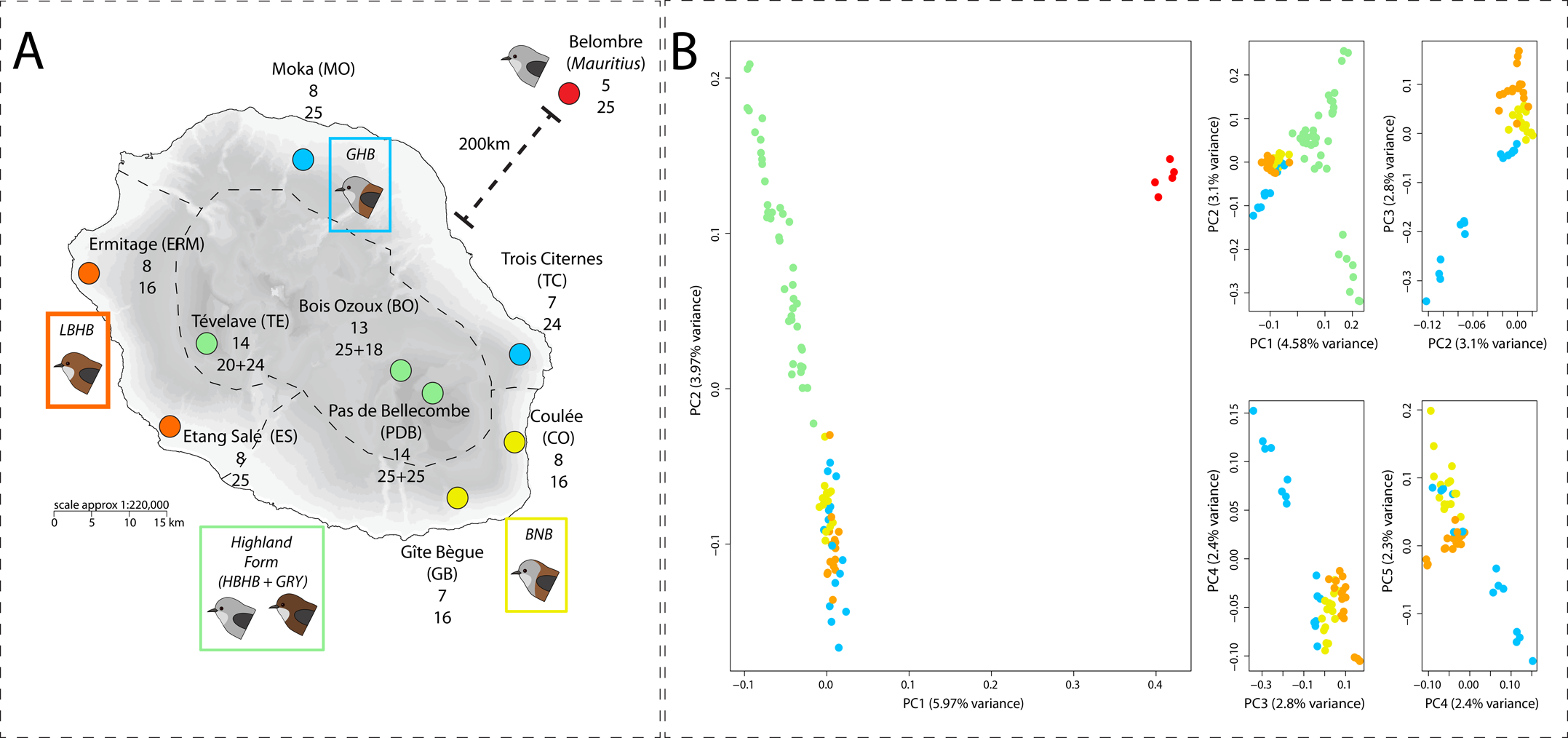
Map of localities and description of population structure using Principal Components Analysis (PCA) on autosomal GBS data. A: For each locality, the sample sizes for the individual GBS dataset is followed by the sample size for each pool. In the three populations found at high elevation, the size of pools is given for grey + brown individuals. B: PCA results including Mauritius (left panel) and without Mauritius (right panels). Points corresponding to high elevation individuals (light green) were removed on the last three panels for better clarity.

Previous studies on this system using microsatellite loci have revealed extremely low natal dispersal rates as indicated by marked genetic structure between populations of the same form (Bertrand et al. 2014), with a likely role of weak dispersal abilities promoting isolation by distance. The island of Réunion displays a steep altitudinal gradient that is associated with strong divergent selection on phenotypes and marked genetic structure for autosomal microsatellites which is consistent with isolation by ecology between forms at high and low elevation (Bertrand et al. 2016). In contrast, plumage forms at low altitude, separated by narrow contact zones at river beds and lava flows, showed no association with neutral genetic structure or major environmental features (Delahaie et al. 2017). Although the autosomal markers used could not provide information on sex-linked loci, this pattern suggests that while the sharp ecological transition between lowlands and highlands could drive differentiation at many autosomal loci through local adaptation, phenotypic divergence in the lowlands involves either fewer loci, or loci concentrated in a narrower genomic region not covered by microsatellites.

In this work, we aim at identifying the genomic variation associated with phenotypic differentiation between forms depending on ecological (low v. high elevation) and morphological divergence (conspicuous variation in plumage colour in the absence of abrupt ecological transition) and determining whether divergence peaks are found on autosomes or sex chromosomes. We used individual genotyping by sequencing (GBS) (Elshire et al. 2011) to characterize the amount of divergence between forms and confirm extensive gene flow between the two most divergent groups. We further used a pooled RAD-sequencing (Baird et al. 2008) approach that produced a high density of markers to characterize with greater precision the genomic landscape of divergence and assess the extent of differentiation between colour forms. Finally, we use coalescent models to test alternative demographic scenarios for the history of gene flow among forms.

## Material and Methods

### Field sampling

Birds were captured in the wild between 2007 and 2012 on the island of Réunion (55°39’E; 21°00’S) using mist nets. Each individual was weighed, marked with a uniquely numbered aluminum ring, and we collected approximately 10 μL of blood. Blood samples were conserved in Queen’s lysis buffer (Seutin et al. 1991) and stored at −20°C. Individuals were sexed by PCR (Griffiths et al. 1998) in order to infer the number of distinct Z chromosomes included in each genetic pool.

### Genotyping by sequencing (GBS) using individual DNA samples

We performed genotyping by sequencing (Elshire et al. 2011) on 95 individuals from 10 localities that were subsequently used to build the pooled DNA samples (Figure 1). Approximately one microgram of DNA was extracted with a QIAGEN DNeasy Blood & Tissue kit following manufacturer’s instructions and sent to the BRC Genomic Diversity Facility at Cornell University (see Elshire *et al.*, 2011). Three individuals were removed due to the extremely low number of reads obtained (Sup. Table 1). Raw reads were trimmed with Trimmomatic (v0.33). We used the recently assembled *Z. lateralis* genome (Cornetti et al. 2015) to map reads back onto this reference with BWA MEM (v. 0.7.12) (Li and Durbin 2009), instead of creating consensuses directly from data as in (Bourgeois et al. 2013). We then aligned contigs and scaffolds from a congeneric white-eye species *Zosterops lateralis* on the Zebra Finch (*Taeniopygia guttata*) passerine reference genome (version July 2008, assembly WUGSC v.3.2.4) using LASTZ (Schwartz et al. 2003). SNPs were called using freebayes (v0.9.15-1) and filtered with VCFTOOLS (0.1.12b) using the following criteria for autosomal markers: i) a mean sequencing depth between 8 and 100X; ii) a minimal genotype quality of 20; iii) no more than 9 missing genotypes. For Z-linked markers, we first called SNPs on males and females and removed markers displaying more than 3 heterozygous females, allowing for some tolerance since freebayes attempts to balance the count of heterozygotes in a diploid population. SNPs were filtered using the same criteria than autosomes for male individuals. The constraint on sequencing depth and genotype quality was removed in females to consider the fact that a single Z copy is found in these individuals, therefore reducing depth of coverage at Z-linked markers.

The final dataset consisted in 34,951 autosomal markers and 965 Z-linked markers. Recent studies have suggested the existence of a neo sex chromosome in Sylvioidae, consisting of the fusion between ancestral Z and W chromosomes with the first 10 Mb of Zebra Finch’s chromosome 4A (Pala et al. 2012). We therefore excluded this region from analyses and studied it separately.

### Pooled RAD-seq

To identify loci and genomic regions associated with environmental and phenotypic features, we used a paired-end RAD-sequencing protocol, using a dataset partially described previously in which 13 pools of 16-25 individuals representing 10 separate populations (Figure 1, Sup. Table 2) were sequenced (Bourgeois et al. 2013). This protocol was used since it produced a high density of markers along the genome compared to the GBS approach mentioned above, facilitating the detection of outlier genomic regions. Reads were mapped on *Z. lateralis* genome using BWA. PCR duplicates were removed using SAMTOOLS (Li et al. 2009). SNPs were called using Popoolation2 (v1.201) (Kofler et al. 2011), using a minimal allele count of 2 and a minimal depth of 10X in all pools. This resulted in more than 1,104,000 SNPs for autosomes and 42,607 SNPs for the Z chromosome.

### Genetic structure

To test for correspondence between phenotypic and genetic structure, we performed a principal components analysis (PCA (Patterson et al. 2006)) on GBS autosomal markers, using the Bioconductor package SeqVarTools (Huber et al. 2015), excluding markers with a minimal allele frequency below 0.05. We evaluated population structure for both autosomal and Z-linked markers using the software ADMIXTURE (Alexander and Novembre 2009). This software is a fast and efficient tool for estimating individual ancestry coefficient. It does not require any a priori grouping of individuals by locality but requires defining the expected number (K) of clusters to which individuals can be assigned. Importantly, ADMIXTURE allows specifying which scaffolds belong to sexual chromosomes, and corrects for heterogamy between males and females. “Best” values for K were assessed using a cross-validation procedure using default parameters. In this context, cross-validation consists in masking alternatively one fifth of the dataset, then using the remaining dataset to predict the masked genotypes. Predictions are then compared with actual observations to infer prediction errors. This procedure is therefore sensitive to heterogeneity in structure across markers induced by, e.g., selection. Therefore, we present results for all values of K as they may reveal subtle structure harboured by only a subset of markers under selection. To assess the relative proportion of markers contributing to colour forms differentiation (estimated by F_CT_) while taking into account population substructure (F_SC_ and F_ST_), we conducted a locus by locus analysis of molecular variance (AMOVA) in Arlequin v3.5 (Excoffier and Lischer 2010) using as groups either islands, lowland and highlands, or colour forms. Significance was assessed with 1000 permutations.

We assessed relationships between populations from pooled data using POPTREE2 (Takezaki et al. 2010) to compute F_ST_ matrices across populations using allele frequencies. A Neighbor-Joining tree was then estimated from these matrices. We included 20,000 random SNPs with a minor allele count of 2. Branch support was estimated through 1,000 bootstraps. As a supplementary control, we also report the correlation matrix estimated from the variance-covariance matrix obtained by the software BAYPASS (v2.1) (Gautier 2015) using both GBS and pooled data. The variance-covariance matrix reflects covariation of allele frequencies within and between populations. The correlation matrix describing pairwise relatedness between populations was then derived using the R function cov2cor() provided with BAYPASS. The function hierclust() was used to perform hierarchical clustering based on matrix coefficients.

### Demography

To assess whether the observed population structure resulted from divergence with gene flow or from recent isolation with no gene flow and incomplete lineage sorting, we estimated the demographic history of the two main groups separating populations from high and low elevation. We used GBS autosomal markers as they could be filtered with higher stringency than pooled markers and were more likely to follow neutral expectations. SNPs mapping on scaffolds corresponding to the neo-sex chromosome region on 4A were discarded from the analysis (see Results). We used a fixed divergence time of 430,000 years with Mauritius and assumed a generation time of 1 year (Warren et al. 2006) to calibrate parameters estimation and provide results in demographic units. This estimate is likely to be conservative, independent analyses (Milá et al. 2010; Bourgeois 2013) having suggested a more recent splitting time (around 100,000 years ago). We compared three distinct models of Isolation with Migration, one with no gene flow between the highlands and lowlands, one allowing constant and asymmetric gene flow, and a last model where gene flow could vary at some time in the past between present time and the split between lowlands and highlands. Population sizes could vary at each splitting time and each group was assigned a specific effective population size. Parameters were estimated from the joint site frequency spectrum (SFS) using the likelihood approach implemented in fastsimcoal2.5 (Excoffier et al. 2013). Parameters with the highest likelihood were obtained after 20 cycles of the algorithm, starting with 50,000 coalescent simulations per cycle, and ending with 100,000 simulations. This procedure was replicated 50 times and the set of parameters with the highest final likelihood was retained as the best point estimate. The likelihood estimated by fastsimcoal2.5 is a composite likelihood, which can be biased by covariance between close markers (Excoffier et al. 2013). To properly compare likelihoods and limit the effects of linkage, we first used a thinned dataset with SNPs separated by at least 10,000 bp. We then used the complete dataset for parameter estimation. We estimated 95% confidence intervals (CI) using a non-parametric bootstrap procedure, bootstrapping the observed SFS 100 times using Arlequin v3.5 (Excoffier and Lischer 2010) and repeating the parameter estimation procedure on these datasets, using 10 replicates per bootstrap run to limit computation time.

### Selection and environmental association

To detect loci displaying a significant association with colouration and environment, we performed an association analysis on the pooled RAD-seq data using the software BAYPASS (Gautier 2015). Divergence at each locus was characterized using *X*^*T*^ *X* statistics, which is a measure of adaptive differentiation corrected for population structure and demography. We also computed the empirical Bayesian p-value (eBPis) to determine the level of association of each SNP with colour forms and elevation. Specifically, we contrasted independently Mauritius, GHB, BNB and HBH pools to all other populations. We considered elevation as a continuous variable. BAYPASS was run using default parameters under the core model. We only included SNPs with a minor allele count of 10 to reduce computation time and ran separate analyses on Z chromosome and autosomes to account for their distinct patterns of differentiation and allele counts.

### GO enrichment analysis

To gain insight into the putative selective pressures acting on *Zosterops borbonicus* populations, we performed a Gene Ontology (GO) enrichment analysis, selecting for each association test SNPs in the top 1% for both eBPis and *X*^*T*^ *X*, analyzing separately Z and autosomes. Gene annotations within 100 kb windows flanking selected SNPs were extracted using the zebra finch reference. We adjusted the gene universe by removing zebra finch genes not mapping on the *Zosterops lateralis* genome. GO enrichment analysis was performed using the package topGO in R (Alexa and Rahnenfuhrer, 2016), testing for significant enrichment using a Fisher’s test for overrepresentation. We present results for GO terms associated with biological processes.

## Results

### Genetic structure and relationships among forms

We first assessed whether colour forms could be distinguished based on the genomic data available for individuals (GBS). A principal components analysis on autosomal allele frequencies revealed a clear distinction between Mauritius and Réunion on the first axis (Figure 1B), as well as a distinction between localities from high and low elevation on the second axis. When excluding Mauritius, the main distinction remained between localities from high and low elevation. Further principal components did not reveal strong clustering of color forms from low elevation (Figure 1B). This pattern was further confirmed by the ADMIXTURE analysis, where localities from high and low elevation were systematically clustered together for both autosomal and Z-linked markers (Figure 2). Cross-validation procedure gave K=2 and K=7 as best models for autosomal and Z-linked markers respectively (Figure S1). Given the subtle genetic structure, we present results for values of K ranking from 2 to 7.

**Figure 2.**
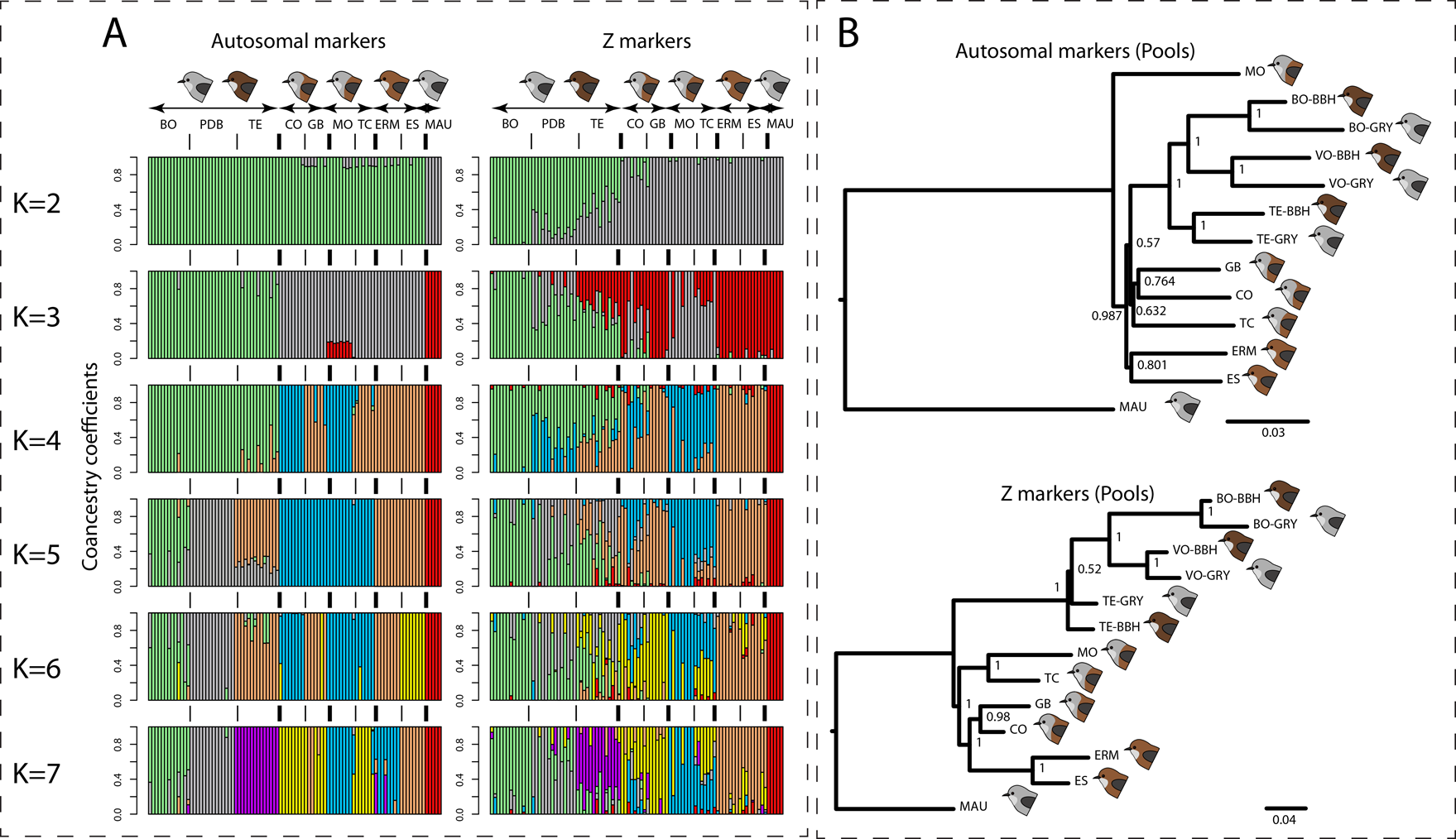
A: Co-ancestry coefficients obtained from ADMIXTURE for K ranging from 2 to 7. Population codes as in Figure 1. Separate analyses were run on autosomal and Z markers. Bold lines and thin vertical lines indicate limits between forms and populations respectively. B: POPTREE2 analysis obtained from 20,000 markers randomly sampled from pooled data. Branch support was obtained from 1,000 bootstraps.

Clustering was consistent with the PCA, with a clear distinction between localities from high and low elevations at K=2 and 3 for Z-linked and autosomal markers respectively (Figure 2A). For autosomal markers, higher values of K highlighted a structure consistent with sampling sites, but structure according to colour forms was more elusive. For Z-markers, clustering tended to be more consistent with colour forms at low elevation. At K=7, three clusters corresponded to localities at high elevation, three others grouped lowlands localities by colour forms, while a last cluster included Mauritian individuals (Figure 2A). The CO and TC localities displayed more signals of mixed ancestry, probably due to high gene flow between those two localities.

Recent studies revealed neo-sex chromosomes in Sylvioidae, a clade to which Zosteropidae belong (Pala et al. 2012). There is however no direct evidence for their existence in this family. These neo-sex chromosomes emerged from the fusion of Z and W chromosomes with the first 10 Mb of chromosome 4A. We confirmed their existence in the Grey White-Eye by extracting GBS markers mapping this region. All these markers were linked to both Z and W chromosomes, as showed by a PCA on allele frequencies (Figure S2). Females (ZW) displayed a strong excess of heterozygous markers, contrary to males (ZZ), due to divergence between neo-Z and neo-W chromosomes. We further investigated population structure in males only, using a set of combined Z and 4A markers. ADMIXTURE analysis tended to better discriminate colour forms with this set of markers (Figure S3).

The same pattern was observed with the POPTREE2 analysis on the pooled dataset. The autosomal topology supported a grouping of localities from high elevation together and supported a grouping of localities belonging to BNB and LBHB forms. There was no support for grouping together localities from the GHB form. Topology based on Z markers provided a good support for a grouping of localities by form (Figure 2B).

We further investigated population structure by estimating the variance-covariance matrix obtained from allele frequencies for both pooled and GBS data using BAYPASS (Gautier 2015). Localities from high elevation were systematically found clustering together, a pattern consistent with previous analyses (Figure S4). Again, both analyses on Z-linked markers for GBS and pooled data revealed a closer relationship between populations from the same colour form when compared to autosomal markers. BHB and BNB forms clustered together, with the GHB form diverging first within the low-elevation group.

To estimate quantitatively the proportion of the genome discriminating among color forms while taking into account population structure within forms, we performed a Molecular analyses of variance (AMOVAs) on GBS data. This analysis confirmed the previous patterns, with a proportion of variance explained by colour forms or elevation not higher than 1.6% for autosomal markers (Table 1). The strongest differentiation was observed between localities from low and high elevation and between Mauritius and Réunion populations. For Z-linked markers however, the proportion of variance explained by colour forms and elevation was more substantial, ranging from 3.4 to 12.6% (Table 1). Differentiation between Mauritius and Réunion was in the same range as nuclear markers.

**Table 1:**
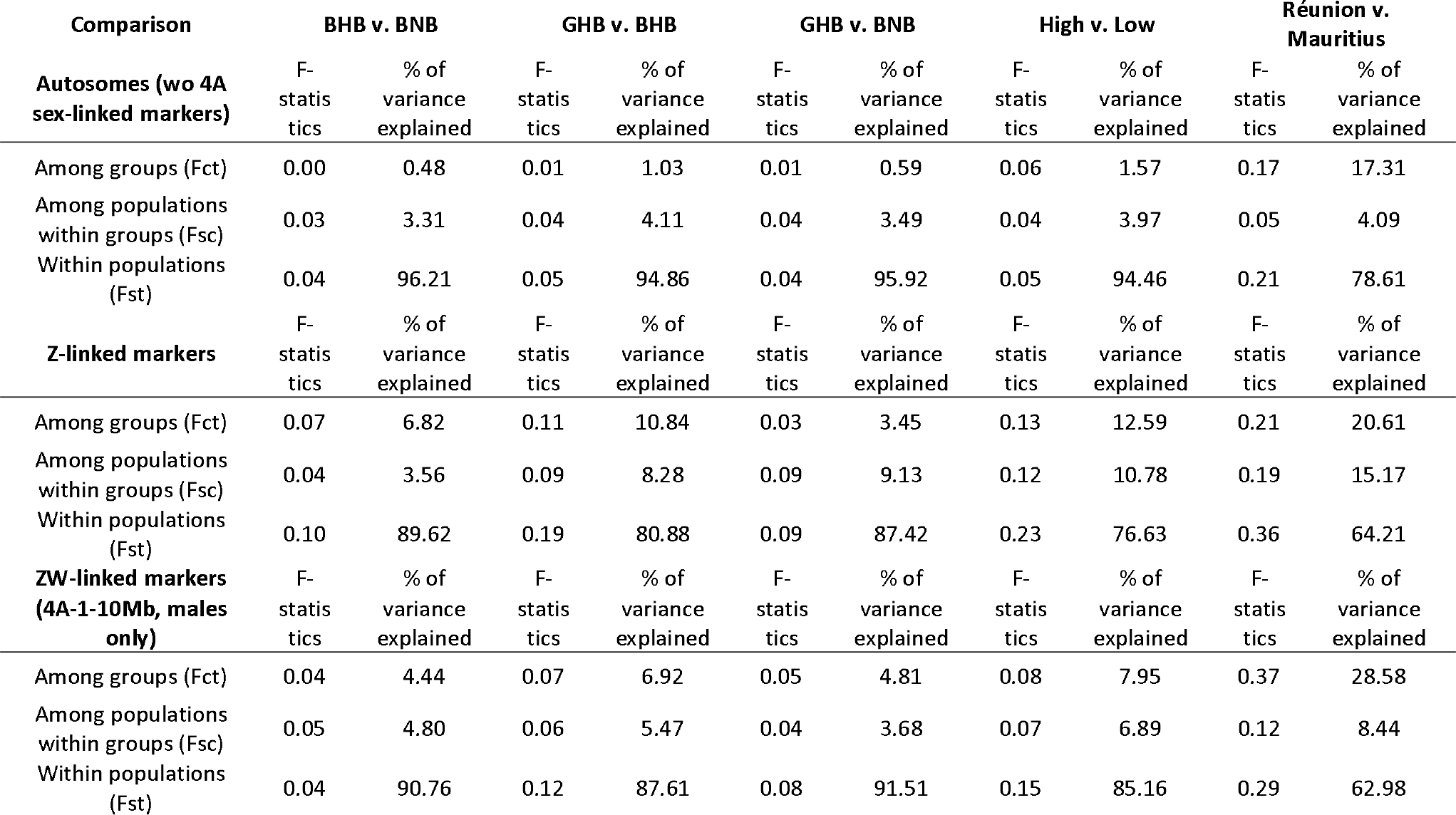
Pairwise AMOVAs comparing forms within Reunion. All values are significant (1000 permutations).

### Genome scan for association and selection analysis

We used BAYPASS (Gautier 2015) on pooled data to retrieve markers displaying high levels of differentiation (*X*^*T*^*X*) and association (empirical Bayesian p-values; eBPis) with 5 different features (altitude, being GHB, being BNB, being BHB, being Mauritian). We examined Z and autosomal markers separately due to their distinct demographic histories. Results revealed a strong association of Z-linked markers with colour phenotypes and altitude (Figure 4). SNPs discriminating Mauritius from other populations were found distributed uniformly along the genome. Most of the peaks displaying a large *X*^*T*^*X* were also found associated with altitude. Strikingly, the clearest peaks on chromosomes 2, 3 and 5 covered large genomic regions, spanning several hundreds of kilobases. The sex-linked region on chromosome 4A was the clearest outlier. Since the neo-W chromosome is highly divergent and found in all females, an excess of variants with frequencies correlated to the proportion of females in the pool is expected. This may lead to high differentiation between pools with different sex ratios. Despite this, the strong association with altitude and colour on chromosome 4A is genuine since the expected proportion of divergent W-linked alleles in each pool was not correlated with those variables in our experiment (Figure S5), making confounding effects unlikely.

**Figure 4.**
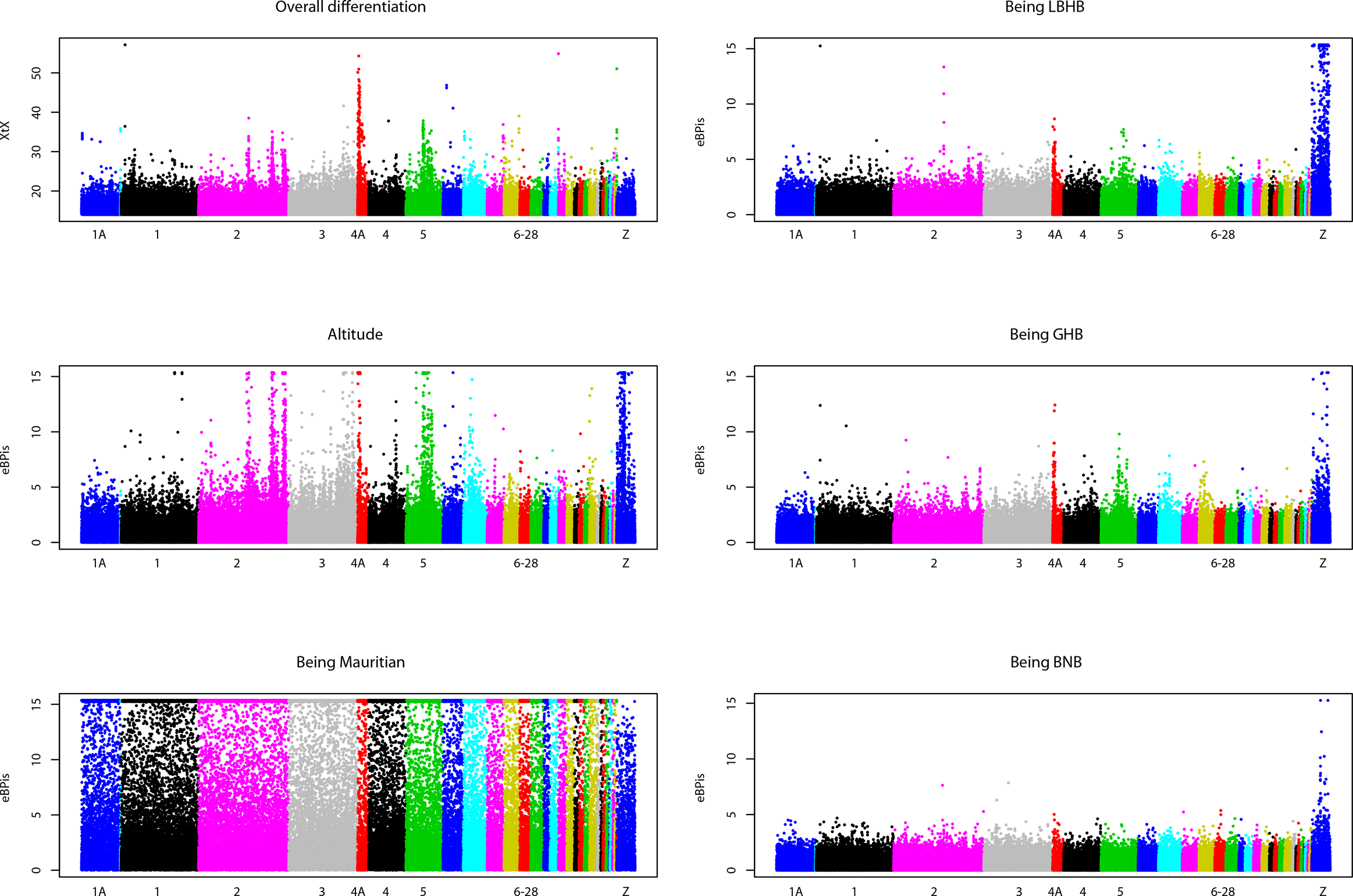
Manhattan plot of *X*^*T*^*X* and Bayesian empirical P-values (eBPis) for association with altitude and phenotype. Note that *X*^*T*^*X* are not directly comparable for autosomes and Z chromosomes as they were analyzed separately and display different demographic histories (Figure 3).

To assess the possible role of these genomic regions in adaptation, we performed a Gene Ontology (GO) enrichment analysis using the zebra finch (*Taeniopygia guttata*) annotation (Supplementary tables 3 to 6). Genes found in regions associated with altitude displayed enrichment for GO terms linked to development, body growth and hemoglobin synthesis (GO:0005833: hemoglobin complex, 3 genes found over 4 in total, P=0.0031). For parapatric forms, we found genes associated with immune response, reproduction, and development.

To identify candidates for colour variation in parapatric forms, we screened the genes found in outlier regions for GO terms linked to melanin synthesis and metabolism (GO:0042438, GO:0046150, GO:0006582). *TYRP1* (*tyrosinase-related protein 1*), located on the Z chromosome, was the only gene with these annotations that was systematically found associated with BNB, GHB and BHB colour forms. Another gene, *WNT5A*, was found associated only with the GHB form (Zhang et al. 2013).

### Demographic history

To confirm the existence of extensive gene flow between forms and rule out incomplete lineage sorting as an explanation for the generally low differentiation, we performed model comparison under the likelihood framework developed in fastsimcoal2.5 (Excoffier et al. 2013) using autosomal GBS data. We focused on the split between populations at high and low elevations, since it was the clearest among all analyses and probably the oldest. We compared three nested models (Figure 3), allowing for no gene flow, constant gene flow, and variable gene flow after the split between populations. The highest likelihoods were found for the model allowing variation in gene flow, while the model with no gene flow could be clearly rejected (ΔAIC = 599.65). Assuming a conservative divergence time with Mauritius of 430,000 years, parameters estimates suggested an increase in gene flow from high altitude to low altitude populations in the last 65,000 years (Table 2) after an initial split 275,000 years ago. Overall, point estimates of effective migration rates (2Nm) were high (2N_High_ m_Low→High_ rising from 0.008 to 9.130 gene copies/generation, 2N_Low_ m_Higi→Low_ rising from 12.0 to 20.4), consistent with homogenization of genomes through recent introgression from low elevation populations into those at high elevation.

**Figure 3.**
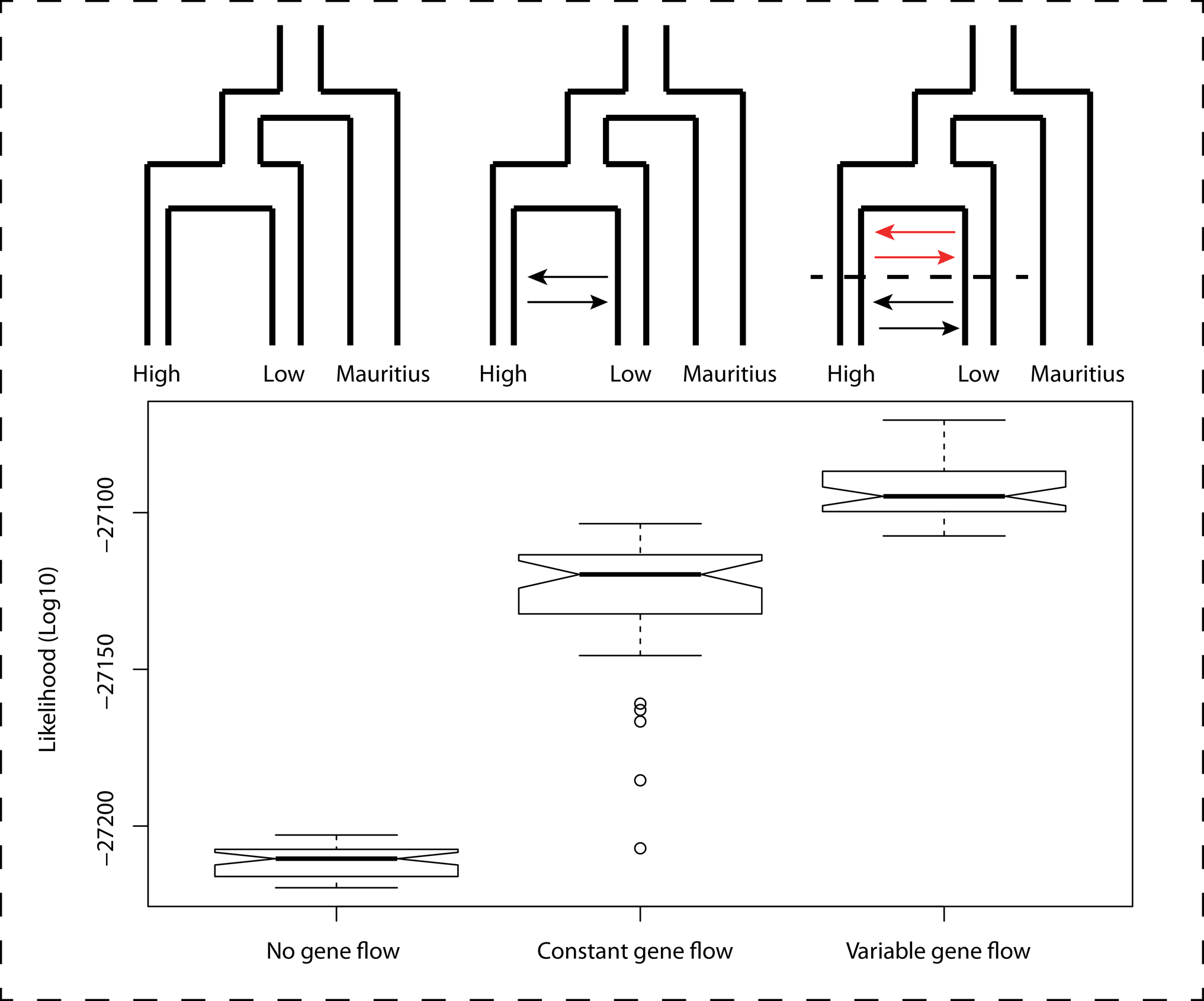
Coalescent tests of three demographic models and their likelihoods across 50 replicates for autosomal markers (GBS data).

**Table 2:**
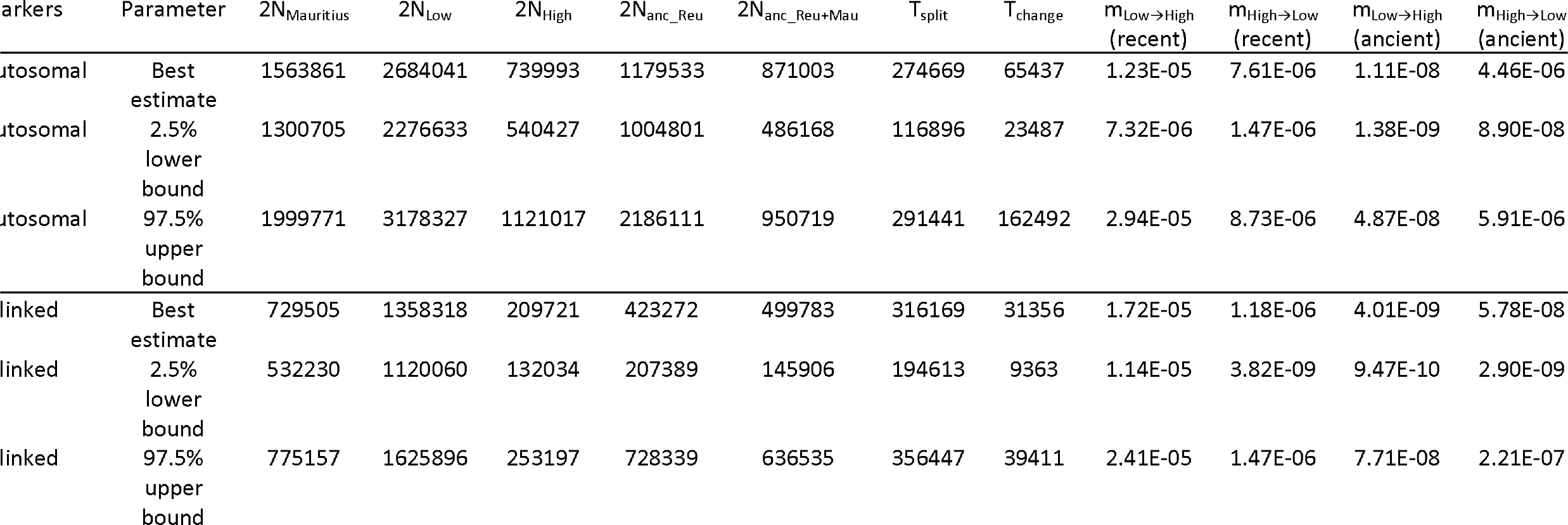
Parameter estimates for a model with variable migration rates. Population sizes are haploid sizes (with N the number of diploid individuals).

Genes that are involved in sexual isolation between populations are expected to resist gene flow due to counterselection of maladapted or incompatible alleles. This should result in an increased variability at these loci when compared to genomic background (Cruickshank and Hahn 2014). We tested this by estimating parameters from the same best-supported model as autosomes on all Z-linked markers. As expected given its haplodiploidy, effective population sizes estimates were lower for this set of markers than for autosomes (Table 2). Estimates of gene flow were also historically lower than for autosomes. This combination of lower population sizes and lower gene flow is expected to lead to increased divergence at Z-linked loci, in accordance with the stronger differentiation observed at these markers in AMOVAs and other tests.

## Discussion

### Genetic structure and genomic islands of differentiation

Our results quantify the relative importance of autosomal and sex-linked genetic variation underlying early speciation in the Réunion Grey White-Eye. We confirm previous findings based on microsatellites data about the existence of a fine-grained population structure, with low but significant F_ST_ between localities (Bertrand et al. 2014), and a clear distinction between populations from low and high elevation (Bertrand et al. 2016). A striking result is the clear contrast between autosomes and the Z chromosome, the latter discriminating more clearly colour forms in parapatry and systematically displaying markers associated with phenotypes and ecology. Demographic analyses suggest that gene flow was never fully interrupted between populations at high and low elevation but may have increased in the recent times. This suggests a role for partial isolation having facilitated initial divergence and promoted local adaptation (Kawecki and Ebert 2004).

### Local adaptation and incompatibilities

Introgression is often found reduced on Z chromosomes, which play an important role in the emergence of reproductive isolation (Carling and Brumfield 2008; Qvarnström and Bailey 2009; Sætre and Sæether 2010). This is in line with our findings, with an excess of highly differentiated markers on the Z chromosome that display evidence for resisting gene flow, a pattern that is also consistent with Haldane’s rule (e.g. Carling and Brumfield, 2008). In that case, mechanisms that can lead to the emergence of highly differentiated loci are intrinsic incompatibilities and post-zygotic barriers. Importantly, the Z chromosome was systematically found associated with both elevation and colour in parapatric forms, while autosomal markers were mostly associated with elevation. This suggests a primary role for reproductive isolation driving differentiation in the Grey White-Eye, while adaptation to environmental conditions seems to play a role mostly along the altitudinal gradient, driving differentiation at multiple loci across autosomes.

GO enrichment analyses suggest a role for adaptation to environment in shaping genetic differentiation in *Zosterops borbonicus*. Genes found in regions associated to altitude displayed an enrichment of genes involved in development and body growth, and included the cluster of hemoglobin subunits A, B, and Z on chromosome 14. The function of these genes is consistent with biological expectations, given the wide altitudinal range (from 0 to 2,500 m) occupied by Grey White Eyes. Previous works have already highlighted the increase in body mass and size-related traits with altitude in the Grey White Eye, consistent with Bergmann’s rule (Bertrand et al. 2016). Altitudinal variation is associated with a strong gradient in mean temperatures, precipitations and oxygen levels, all factors that exert a selective pressure on body mass and oxygen intake.

For parapatric forms found at low elevations, there was an enrichment in genes involved in immune response and reproduction. These functions make biological sense since bird populations in Réunion are infected by communities of parasites such as the malaria vector that are spatially structured (Cornuault et al. 2013), and could therefore exert disruptive selection at immune genes.

Large chromosomal rearrangements are powerful drivers of differentiation, since they prevent recombination between several consecutive genes, facilitating the maintenance of divergent allele combinations between populations. A famous example of adaptive inversion facilitating the maintenance of colour forms and species has been reported in *Heliconius* butterflies (Joron et al. 2011), and these rearrangements have been predicted to be favored by isolation and secondary contact (Feder et al. 2011). Autosomal regions associated with colour forms and altitude sometimes spanned more than 1 Mb, too long to be explained by a single selective sweep. This suggests a possible role for large scale rearrangements or regions of low recombination as a substrate for divergence in *Zosterops borbonicus*. Our results remind at a much smaller spatial scale what has been previously observed in *Ficedula* flycatchers, with high differentiation on the Z chromosome and possibly large-scale genomic rearrangements (Ellegren et al. 2012).

Demographic analyses suggest that diversification in *Zosterops borbonicus* has taken place in a context of high, increasing gene flow between lowland and highland forms, which can have facilitated coupling (Bierne et al. 2011) between incompatibility loci and loci linked to local adaptation once populations became more connected. This is remarkable since most studies have so far neglected the possible role of incompatibilities when high, even continuous gene flow connects populations (Kulmuni and Westram 2017). Future studies should focus on the variation in allele frequencies along hybrid zones and test whether loci that are more likely to be involved in local adaptation (such as immune genes or hemoglobin subunits) display changes in frequencies that are as sharp as Z-linked loci, since the latter are more likely involved in pre- and post-zygotic isolation.

### Genetics of colour and assortative mating

*TYRP1*, a well-characterized colour gene in both model species and natural populations (Nadeau et al. 2007; Backström et al. 2010; Delmore et al. 2016), was the only known colour gene systematically found among the regions associated with colour forms in lowlands. This gene had been previously studied using a candidate gene approach, and displayed low polymorphism in lower altitude populations, preventing the detection of any association with phenotype (Bourgeois et al. 2016). Together with previous findings (Bourgeois et al. 2012, 2017), this suggests that colour variation in *Zosterops borbonicus* may be controlled by a set of two loci of major effects. Yet other undescribed colour loci may also be involved, and more detailed studies on birds sampled in hybrid zones may help characterizing the exact association of alleles that produce a given phenotype.

Candidate colour genes in the Grey White Eye are involved in intracellular vesicular transport and pigment synthesis (Bourgeois et al. 2017), and are therefore less likely to display pleiotropic effects as strong as the ones displayed by *ASIP* or *POMC* (Ducrest et al. 2008). This could mean that selection acts on colour itself, and not on another trait controlled by the colour gene, consistent with expectations in the case of sexual selection. Indeed, color forms are distinguished by birds (Cornuault et al. 2015) and colour variation seems to be under strong selection.

It is theoretically possible for assortative mating alone to maintain divergence in species that are sympatric or with high gene flow between them. However, there is a strong expectation that prezygotic preferences emerge to reduce the odds of crosses between individuals harboring incompatible alleles, increasing differentiation (Kirkpatrick 2000; Servedio and Noor 2003; Hall and Kirkpatrick 2006). Such patterns of reinforcement are common in recently diverged bird species for which Z chromosome differentiation is high (Pryke 2010; Sætre and Sæether 2010; Ellegren et al. 2012), though none has ever focused on the case of recent divergence with gene flow at the spatial scale highlighted in this study. More research about the ecology of the Grey White-Eye is needed to quantify the extent of assortative mating, parental imprinting, intrinsic incompatibilities, and how these factors interact in this system (Pryke 2010; Seehausen et al. 2014). Our results suggest an extreme case of divergence with gene flow that would bring valuable insights about the relative order at which pre- and post- zygotic isolation mechanisms occur during speciation. *TYRP1* is located on the Z chromosome, which could facilitate linkage with other incompatible loci involved in reinforcement (Hall and Kirkpatrick 2006) or hybrid fitness (Lemmon and Kirkpatrick 2006). In that case, colour variation could constitute an emerging prezygotic barrier, with hybrid phenotypes signaling maladapted combinations of alleles, promoting reinforcement.

## Supplementary Figures

Figure S1. Cross-validation error plots for ADMIXTURE analyses and a range of K values.

Figure S2. PCA on allele frequencies for markers mapping on the first 10Mb of Zebra Finch’s chromosome 4A.

Figure S3. ADMIXTURE analyses for concatenated Z-linked markers and ZW-linked markers from Zebra Finch’s chromosome 4A in male individuals.

Figure S4. Relationships between populations for pools and GBS datasets inferred from BAYPASS.

Figure S5. Plot of the expected proportion of divergent W copies in RAD-seq pools v. elevation. Points are colored according to forms. Correlation between elevation and proportion of W copies: Spearman’s rho=−0.8852, P-value=0.77. Effect of colour forms on proportion of W copies: Kruskal-Wallis test, P-value=0.58.

### Supplementary tables

**Table S1:**
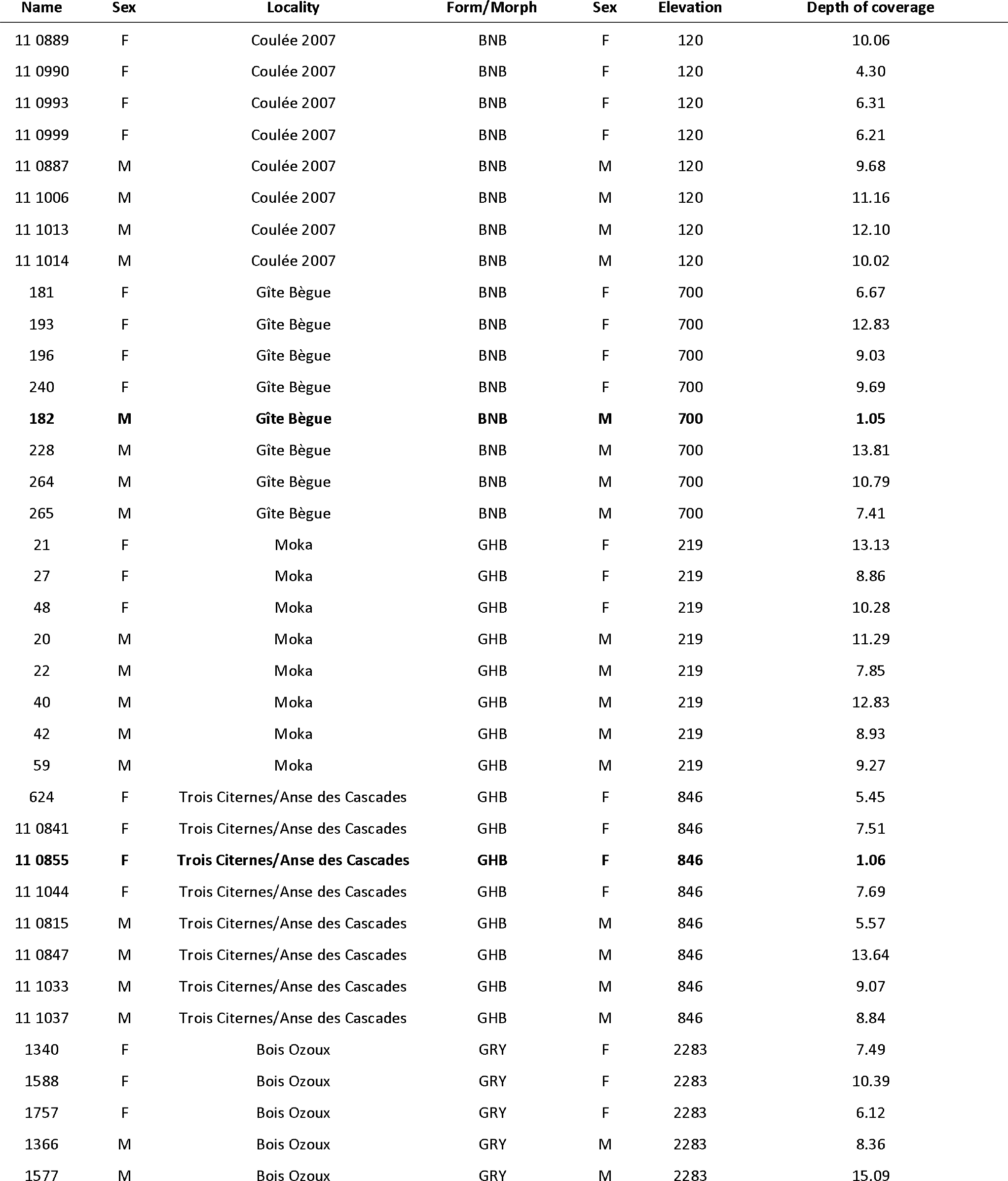
List of individuals included in GBS analyses with their sex, sampling sites and mean depth of coverage before filtering. Excluded individuals are highlighted in bold.

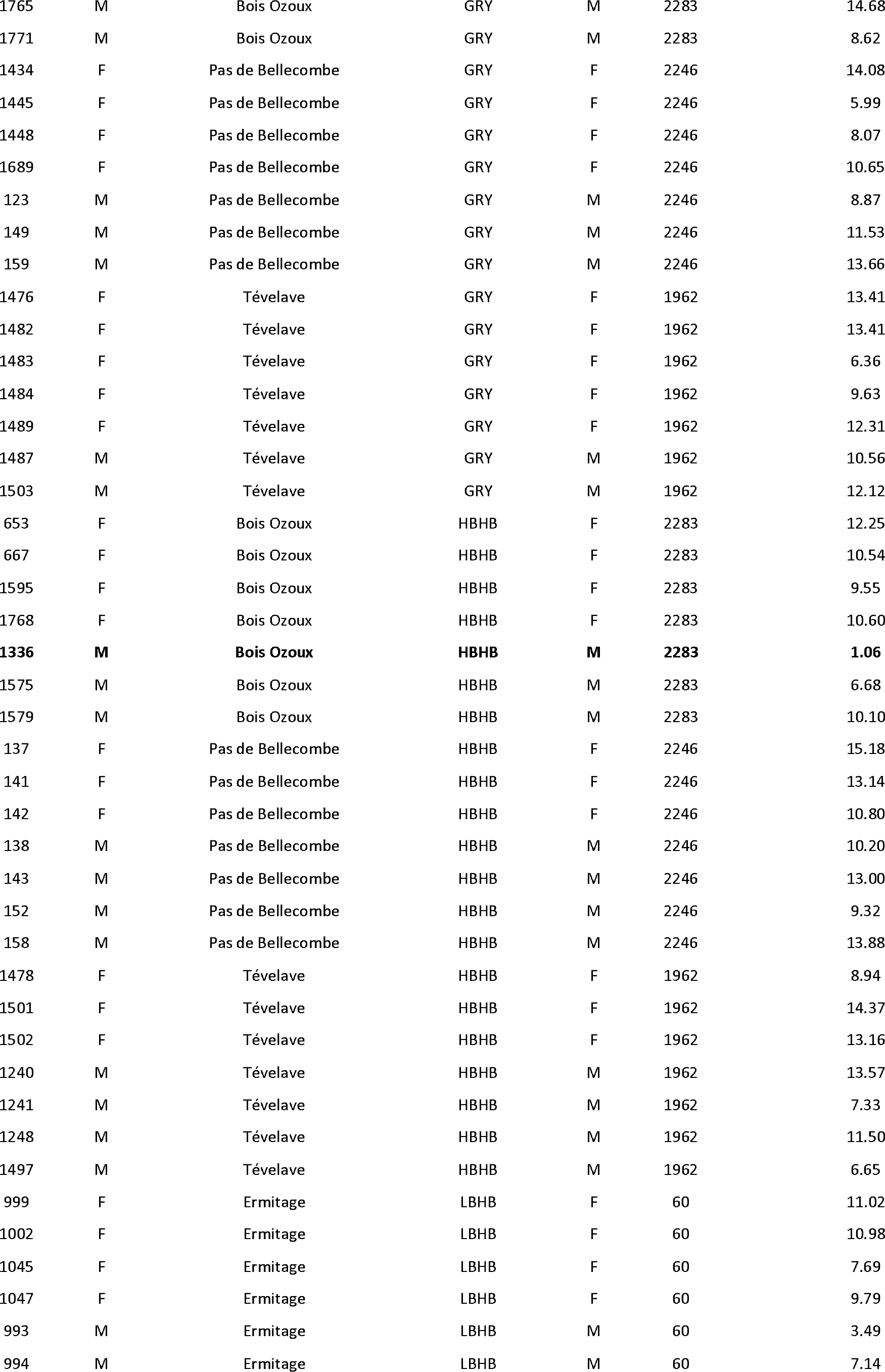

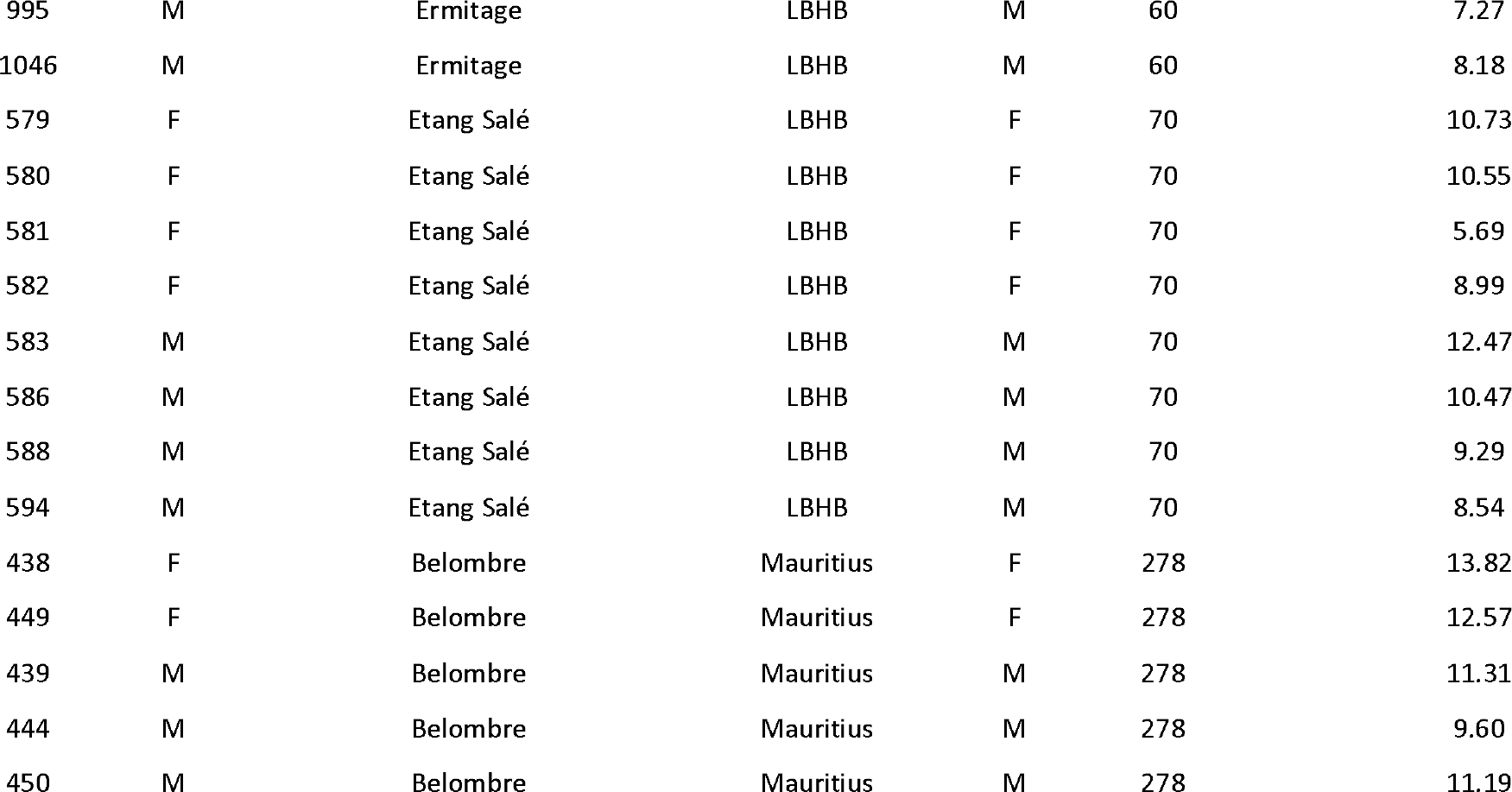

**Table S2:**
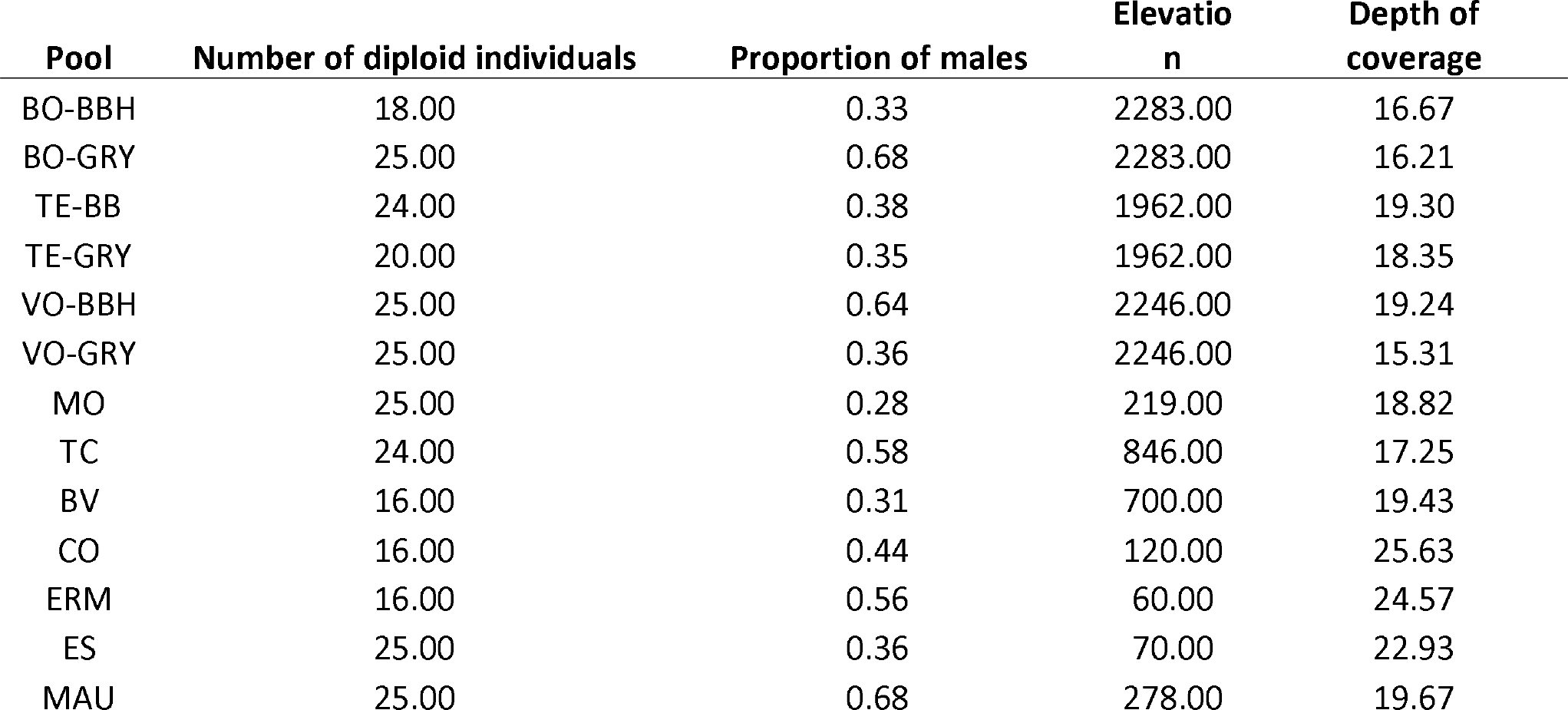
Elevation, mean sequencing depth before filtering and sex-ratio for all RAD-sequencing pools.

**Table S3:**
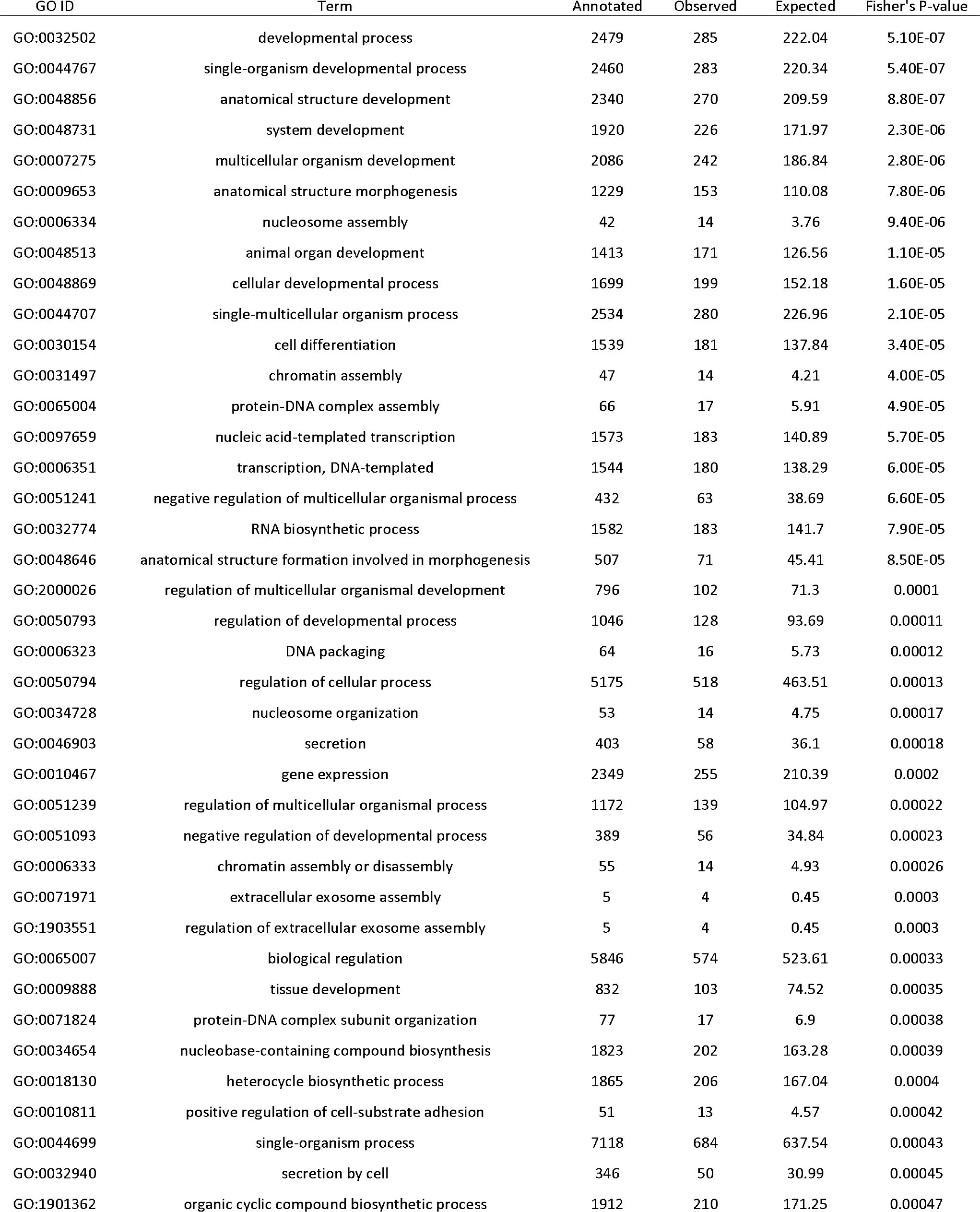
GO terms (biological process) associated with altitude. The 50 most significant terms are shown.

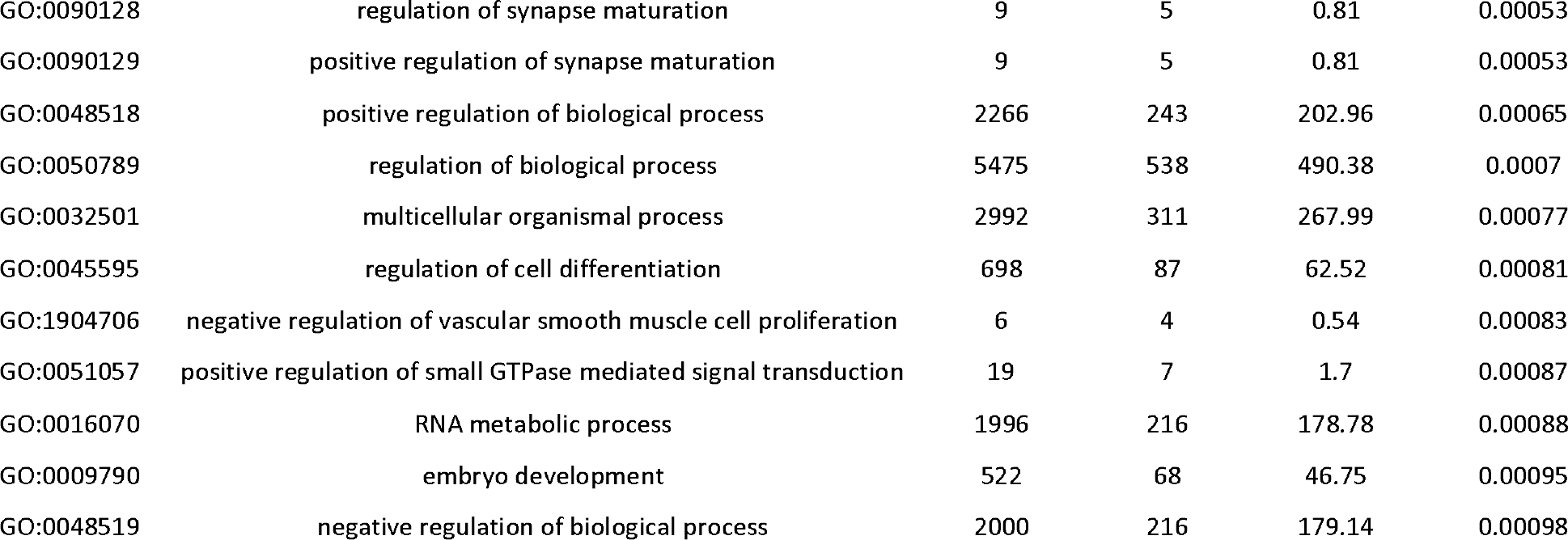

**Table S4:**
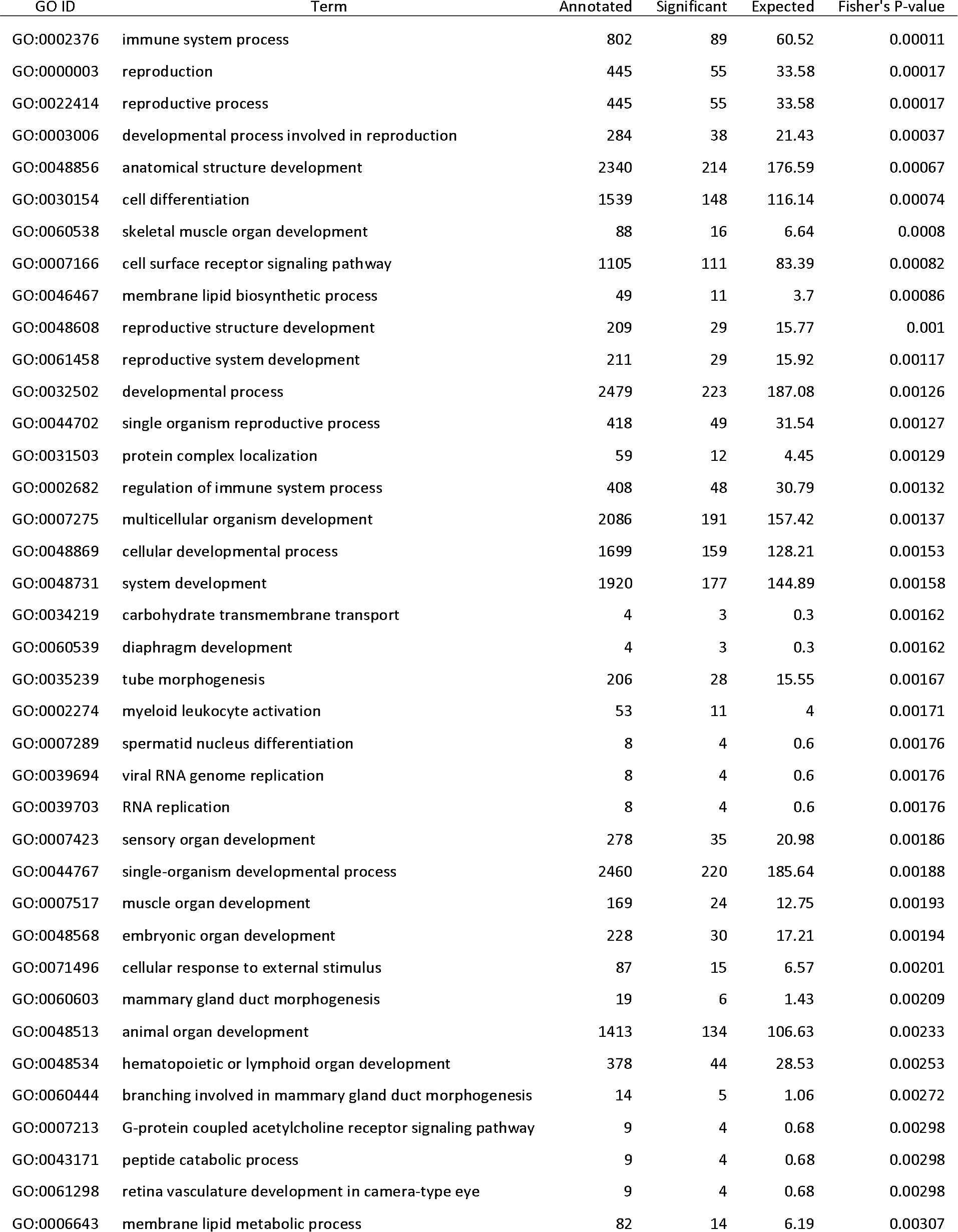
GO terms (biological process) associated with the GHB form. The 50 most significant terms are shown.

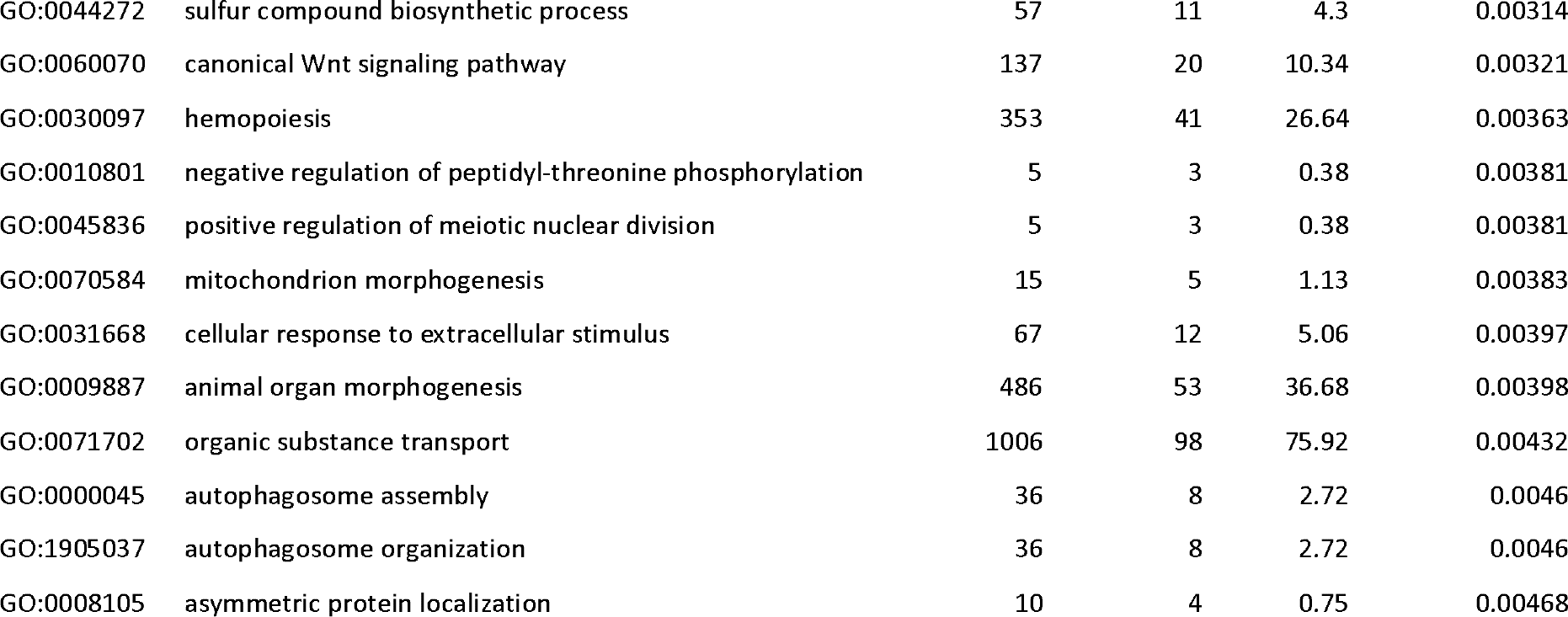

**Table S5:**
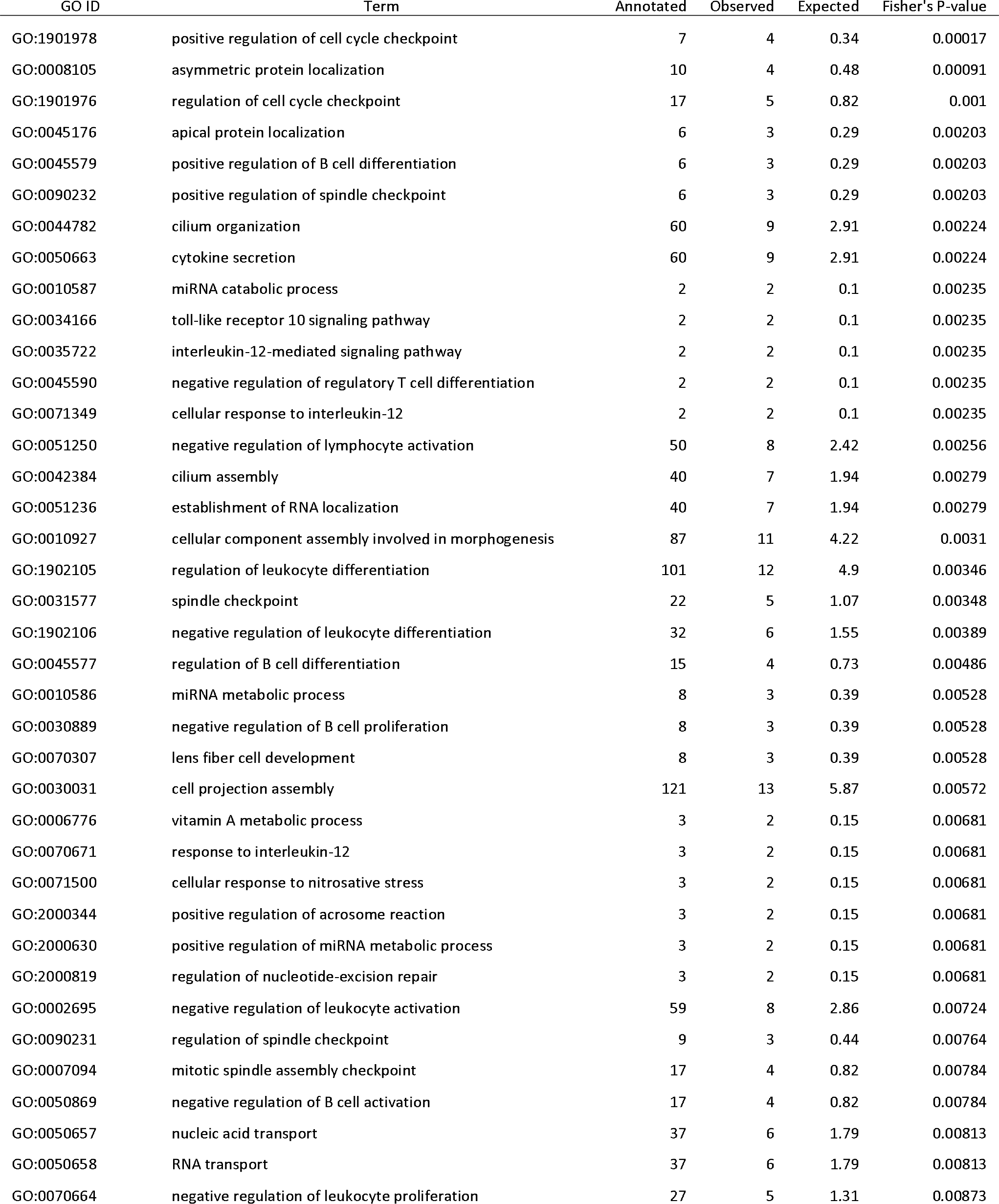
GO terms (biological process) associated with the BNB form. The 50 most significant terms are shown.

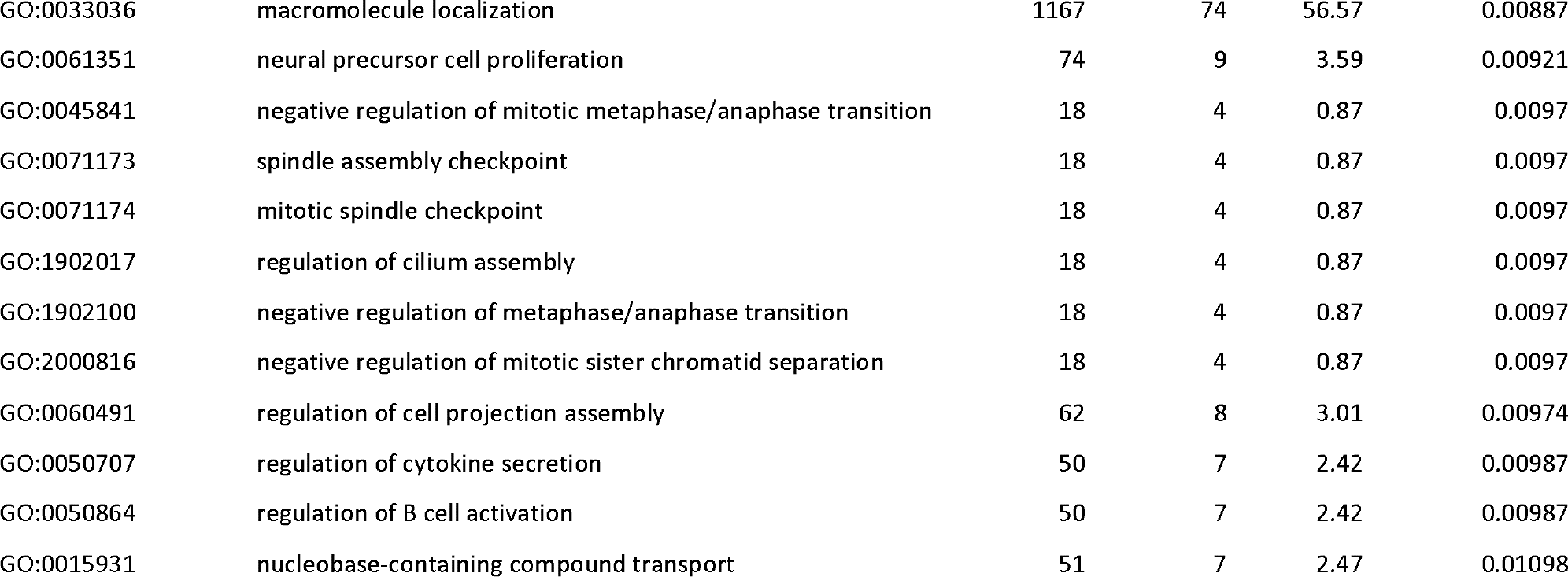

**Table S6:**
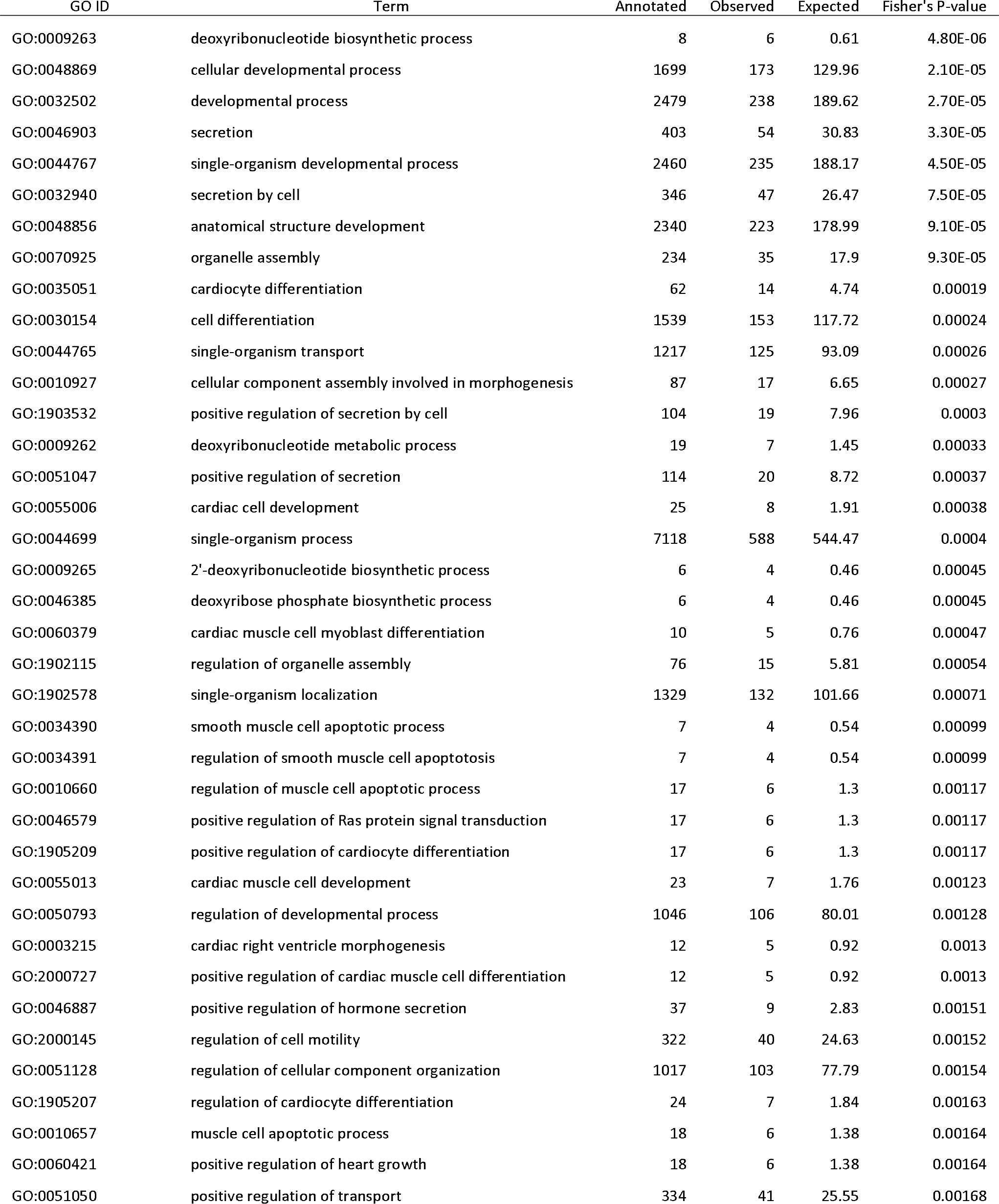
GO terms (biological process) associated with the BHB form. The 50 most significant terms are shown.

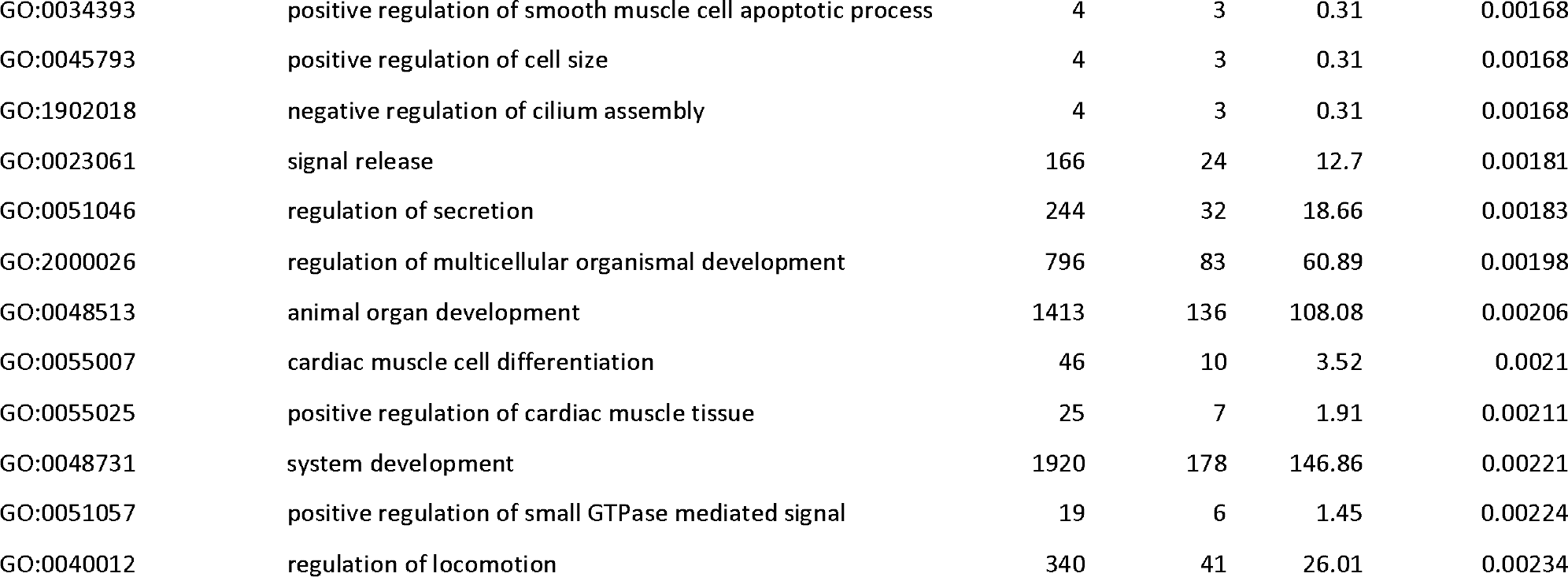

